# Unlocking the Predictive Power of Heterogeneous Data to Build an Operational Dengue Forecasting System

**DOI:** 10.1101/2020.07.08.194019

**Authors:** Carrie Manore, Geoffrey Fairchild, Amanda Ziemann, Nidhi Parikh, Katherine Kempfert, Kaitlyn Martinez, Lauren Castro, David Osthus, Amir Siraj, Jessica Conrad, Nicholas Generous, Sara Del Valle

## Abstract

Predicting an infectious disease can help reduce its impact by advising public health interventions and personal preventive measures. While availability of heterogeneous data streams and sensors such as satellite imagery and the Internet have increased the opportunity to indirectly measure, understand, and predict global dynamics, the data may be prohibitively large and/or require intensive data management while also requiring subject matter experts to properly exploit the data sources (e.g., deriving features from fundamentally different data sets). Few efforts have quantitatively assessed the predictive benefit of novel data streams in comparison to more traditional data sources, especially at fine spatio-temporal resolutions. We have combined multiple traditional and non-traditional data streams (satellite imagery, Internet, weather, census, and clinical surveillance data) and assessed their combined ability to predict dengue in Brazil’s 27 states on a weekly and yearly basis over seven years. For each state, we nowcast dengue based on several time series models, which vary in complexity and inclusion of exogenous data. We also predict yearly cumulative risk by municipality and state. The top-performing model and utility of predictive data varies by state, implying that forecasting and nowcasting efforts in the future may be made more robust by and benefit from the use of multiple data streams and models. One size does not fit all, particularly when considering state-level predictions as opposed to the whole country. Our first-of-its-kind high resolution flexible system for predicting dengue incidence with heterogeneous (and still sometimes sparse) data can be extended to multiple applications and regions.

## Introduction

The emergence of recent deadly epidemics has highlighted the need for anticipatory modeling approaches and tools to improve outbreak preparedness and response ^1, 2^. Specifically, mosquito-borne diseases such as Zika and chikungunya have been on the rise but forecasting their spread is challenging due to the lack of near real-time data to inform computational models that can provide decision support ^3^. For example, long-term and consistent measurements of mosquito populations are rare, as is availability of robust reporting of human case counts ^4, 5^. Thus, there is a need for the use of *proxy data* (i.e., alternative, indirect data sources) to derive relevant information that can capture the complex interactions between the host, pathogen, and environment. While such indicators may be imperfect, they can provide information for local conditions in the absence of other granular statistics collected on the ground. Targeted studies at relatively small spatial scales have illustrated the usefulness of proxy data such as weather/climate ^6–8^, demographics ^9^, multispectral satellite imagery ^10, 11^, and social internet data ^12, 13^ to enhance predictions. However, the field is limited by a lack of consistent metrics, the need for broad comparative assessment of data streams, and the need for methods that can work across the complex matrix of eco-regions and human systems where mosquito-borne diseases are endemic.

In response to recent outbreaks, there has been an explosion in the number research papers related to disease nowcasting and forecasting ^14, 15^ as well as on the applicability of data fusion approaches ^16, 17^ to capture various aspects of infectious disease dynamics. Accurate, real-time forecasting can inform public health decision makers to allow for improved response, targeted surveillance, and improved mitigation^18^. Broadly speaking, dengue prediction studies have focused on the Americas ^19^ and Southeast Asia ^20^. Time scales range from predicting categorical risk for an entire transmission season ^21^ to weekly case counts ^19^. Spatial scales are most commonly limited to cities ^22^, small countries such as Singapore ^20^, or specific regions (e.g., county, state) within a country with particularly good human case counts and/or mosquito time series data ^23^. Windows of prediction include nowcasting (necessary since reported case counts often have a lag of 1–4 weeks) ^19^, forecasting weekly cases 4–12 weeks into the future ^20^, and several-months ahead categorical risk forecasting ^21^. Many of these studies have exploited multiple data streams including socioeconomic drivers ^9^, weather data ^6^, satellite imagery ^11^, topography (e.g., altitude), entomological factors ^24^, Internet data (e.g., social media) ^13, 25^, and clinical surveillance data ^19^. However, most of these studies have not evaluated the contribution of each data stream on the forecast, with a few exceptions that have evaluated the contribution of one data source ^12^ or a single type of data stream ^20^. Only one study^26^ developed a systematic approach to evaluate the contribution of multiple data sources (lagged case counts, weather, case counts from the neighboring districts, and socioeconomic data) for the city of Bangkok, and showed that a model that combined all of this information led to the optimal performance.

To address the aforementioned gaps, this paper takes the first step towards creating an operationalizable framework that quantifies the contribution of disparate data streams in predicting dengue in Brazil. We assess the relative utility of heterogeneous data streams (i.e., weather, satellite imagery, demographics, and Google Health Trends) in predicting dengue. No prior work has used all of these data sources together to predict dengue at a state scale or finer resolution. Second, we build models at various spatial (i.e., national, state, and municipality) and temporal (i.e., yearly and weekly) resolutions to account for the highly heterogeneous dengue spread in Brazil (Fig. 1). Finally, we find that different statistical models (or an ensemble of models) and combination of data streams across space and time can accurately predict dengue. We present these results with error metrics comparable to other methods and models. These results help not only identify which data sources are useful for predicting dengue but also identify alternate data sources in the absence of ground truth data. Our approach can provide the foundation for a robust operational system for nowcasting and forecasting dengue across broad geographical regions.

**Figure 1.**
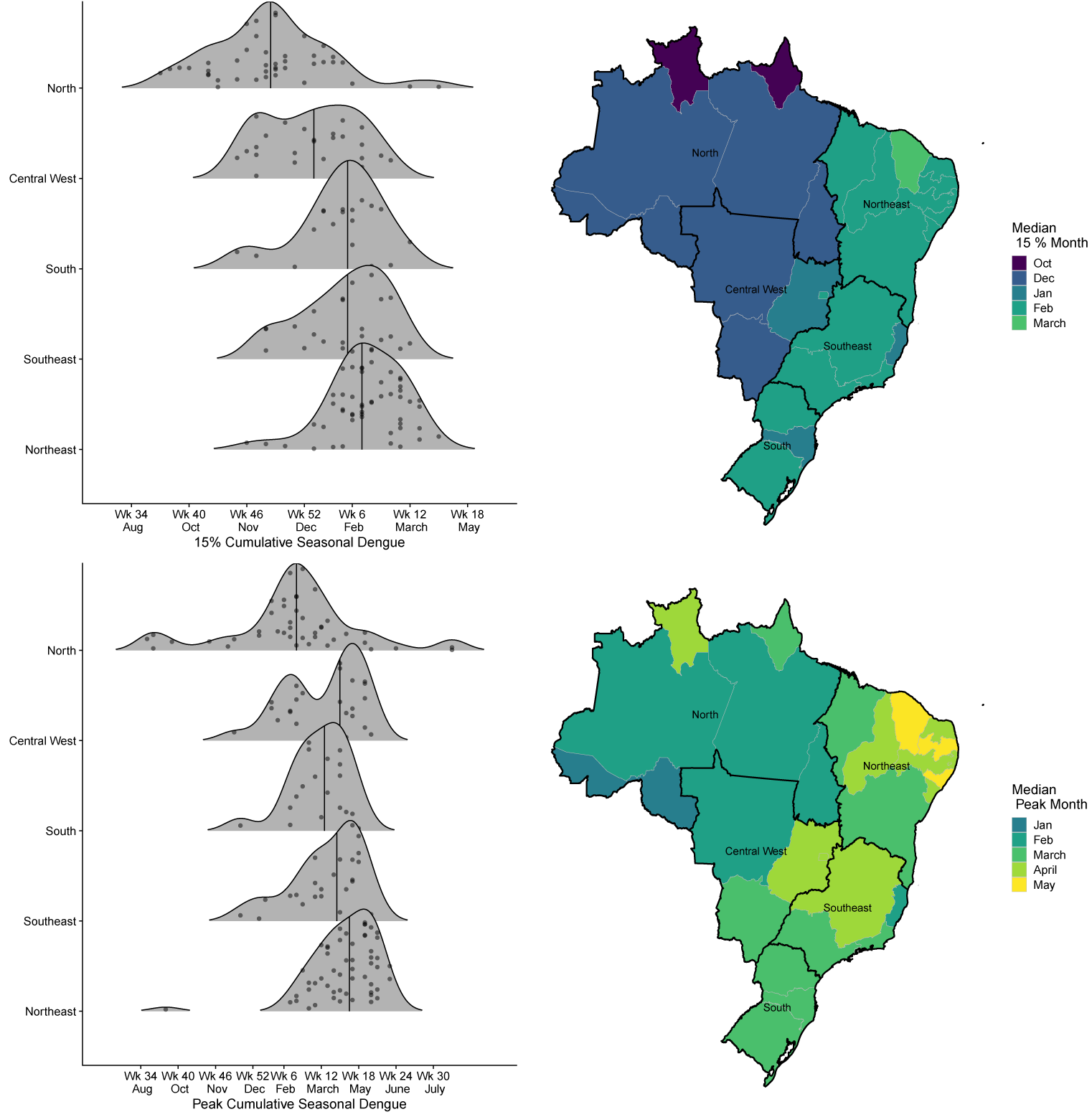
Dengue timing in Brazil, 2010-2016. LEFT: Density plots showing the calendar week when states reach an early warning threshold (15% of seasonal cumulative cases, TOP) and peak seasonal cases (BOTTOM) for each region as defined by the Brazilian government. Each dot represents a state’s value from 2010–2016. The line represents the median timing of each region. RIGHT: Maps showing the month associated with the median calendar week of early warning (TOP) and peak thresholds (BOTTOM) across Brazil.

## Results

We first calculated risk maps of yearly cumulative dengue incidence using demographic data and aggregated satellite imagery data, weather data, and Google Health Trends (GHT) data as exogenous predictors at the municipality and state scales. We then used three base statistical models and their variants with and without exogenous predictors to produce weekly nowcasts of dengue at the state scale: seasonal autoregressive integrated moving average (SARIMA), vector autoregression (VAR), and seasonal trend decomposition based on locally estimated scatterplot smoothing (STL). Considering more than 100 potential predictors, we produced weekly nowcasts (forecasts two weeks ahead of available data) of dengue case counts for every state in Brazil as reported to the Brazil Ministry of Health. The exogenous variables used for time-series prediction were indices derived from satellite imagery, weather, and GHT. We found for yearly and weekly predictions that best model performance required sub-country model building and model selection. The best models state-by-state used input from subsets or occasionally all of our exogenous data streams, as shown in Fig. 2. Accuracy of our models varied across states and was found to be correlated with volatility of dengue case counts data, the total number of dengue cases per year, and the strength of the seasonal signal for each state. Additionally, demographic variables having to do with education and poverty were correlated with the accuracy of our predictions.

**Figure 2.**
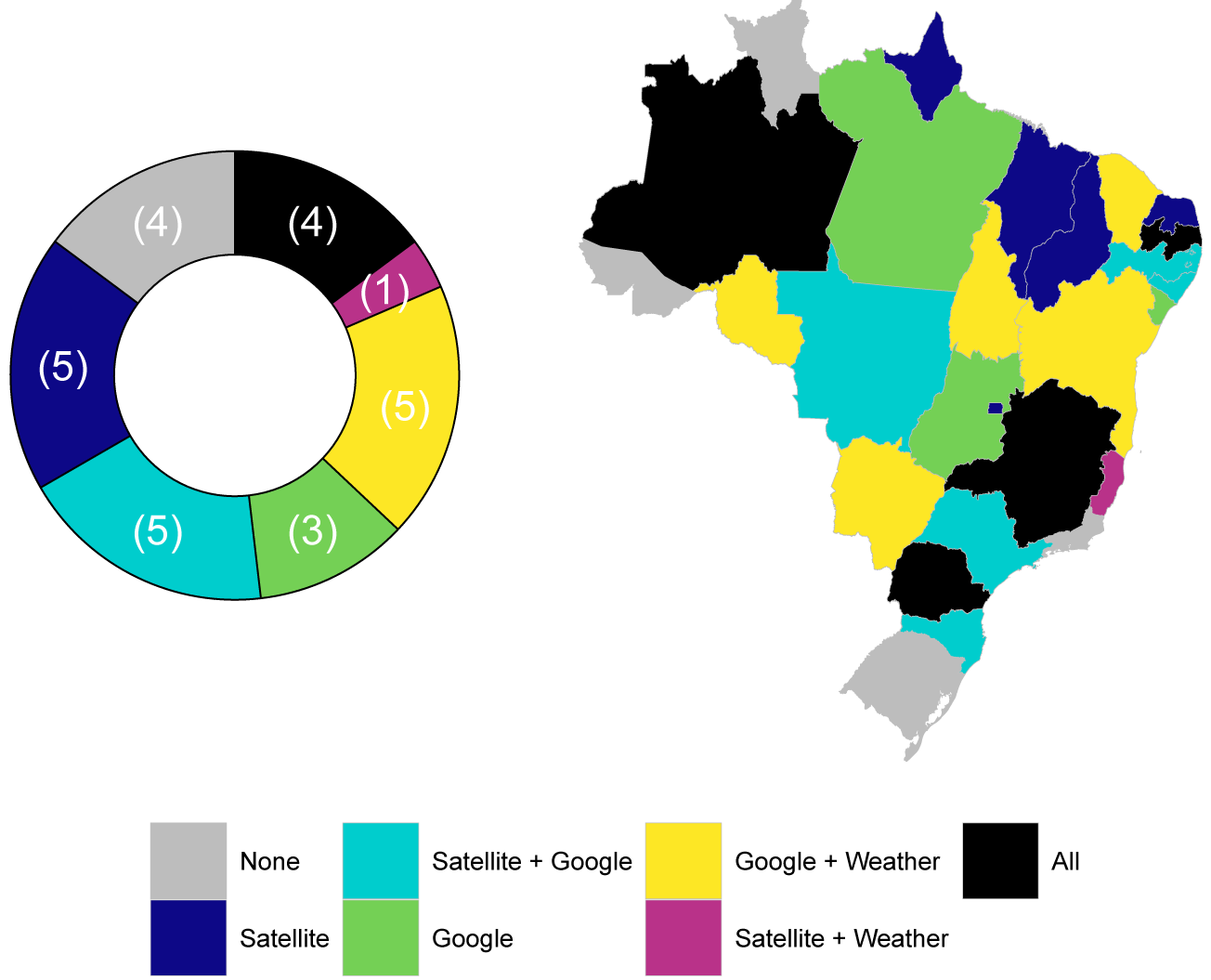
Number of data streams used by the best nowcasting model for each state. LEFT: Among 27 states, four were predicted best with no exogenous data (only past case counts, STL model), while four required all three exogenous data streams for best performance. Eight did best with a single additional data streams and 11 with a combination of two data streams. RIGHT: Exogenous data streams used by the best individual nowcasting model by state.

### Yearly risk maps

Yearly risk by state was more accurate when we built a model for each of the five regions separately rather than the whole country. The risk maps pulled out demographic, satellite imagery, and weather variables as top predictors, where the chosen variables and their relative importance varied by region in Brazil. The five political regions in Brazil correspond roughly to the main ecoregions in Brazil: Amazon rain forest (North), dry forest (Northeast), savanna (Center West), Atlantic forest (Southeast), and subtropical grassland (South) as well as Pantanal wet lands that also occur in Center West. Brazil’s population is highest in the Southeast, Northeast, and South regions, and the number of cases is highest in the Southeast and Northeast where population density and ecological suitability meet. However, incidence, or cases per 100,000 people, varies more widely across regions depending on ecology, population density, and human system characteristics such as poverty and infrastructure.

### Nowcasting weekly incidence

As with yearly incidence, the accuracy of nowcasting improved when we built models by state rather than choosing a best model for the whole country. SARIMA performed better than VAR with exogenous variables and so we are presenting results here for SARIMA and STL. We compare two time series statistical methods that include only past dengue case counts as predictors (STL and SARIMA) as well as two including exogenous data, SARIMA with exogenous variables with PCA or PLS (SARIMAX). Based on three standard error metrics, relative RMSE (RRMSE), relative mean absolute error (RMAE), and R correlation, and a ranking of models with a combination of these three metrics, we found that different statistical methods worked better in different states (see Fig. 4). Roraima, Acre, Espirito Santo, and Rio Grande do Sul performed best with no exogenous variables using a seasonal trend decomposition model (STL). Five states performed best with SARIMAX-PLS, one with SARIMA, and the rest with SARIMAX-PCA. In light of the variation in accuracy among models, we also tested an ensemble model. For a trimmed mean ensemble approach, 25 states achieve Pearson correlation coefficients (between fitted and observed values in the 2015-16 testing window) exceeding 80%; meanwhile, the median value over the 27 states is 91.75%, and the maximum is 96.44%. Associated 95% prediction intervals reach approximately 96% empirical coverage or higher for half the states

**Figure 3.**
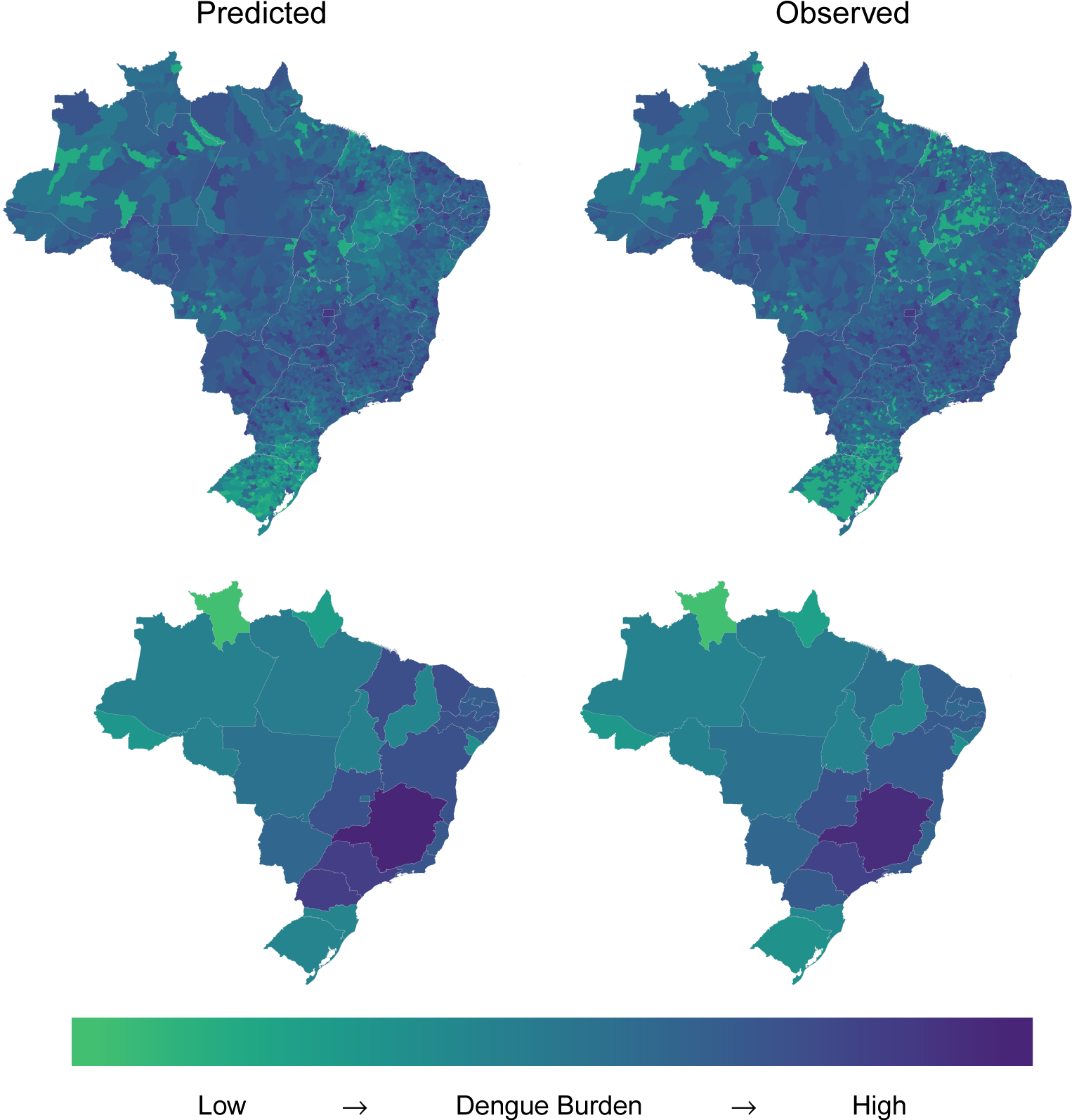
2016 observed and predicted dengue burden. Results from linear models that were trained on 2010–2014 dengue cases at the municipality level and tested on the withheld data from 2015 and 2016. The predictors in the linear model come from yearly summaries of the satellite imagery data, weather data, and the 2010 Brazil census. TOP: maps showing the municipality level results. BOTTOM: maps showing the results aggregated to the state scale.

**Figure 4.**
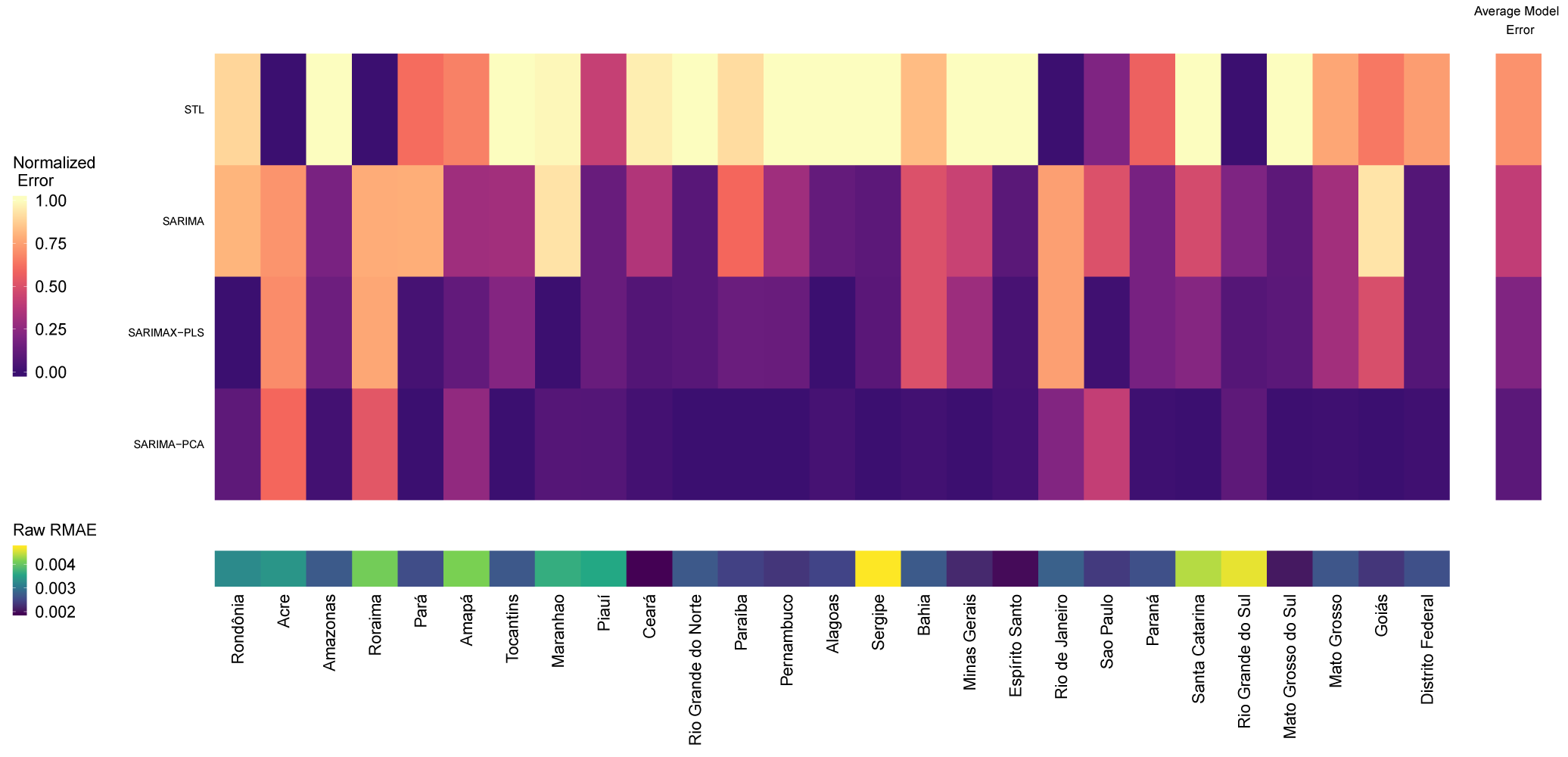
Relative and absolute performance of nowcasting models across Brazil. TOP: Heat map showing the normalized average error (combining RMAE, R, and RRMSE) for each model by state. Purple indicates a better model fit. The “Average Model Error” column shows the average normalized error across all of the states. Different models perform better for different states and SARIMA-PCA performs best overall. BOTTOM: Using the model with the lowest normalized average error for each state, the performance, measured by the raw RMAE values, varies across the states. Dark blue is lower error and yellow is higher error. For our models and data streams, some states are more “predictable” than others.

For most states, adding exogenous variables increased the accuracy of performance of our nowcasting model (Fig. 5). The data streams or combination of data streams that provided the most improvement again varied by state, with combinations of Google search queries and either weather or satellite imagery being the most common (Fig. 2). Four states required all three data streams for best performance (Amazonas, Minas Gerais, Paraiba, and Parana). When aggregated for the year, the predicted and true risk from the nowcasting model agrees well with our yearly risk map prediction and the true number of cases.

**Figure 5.**
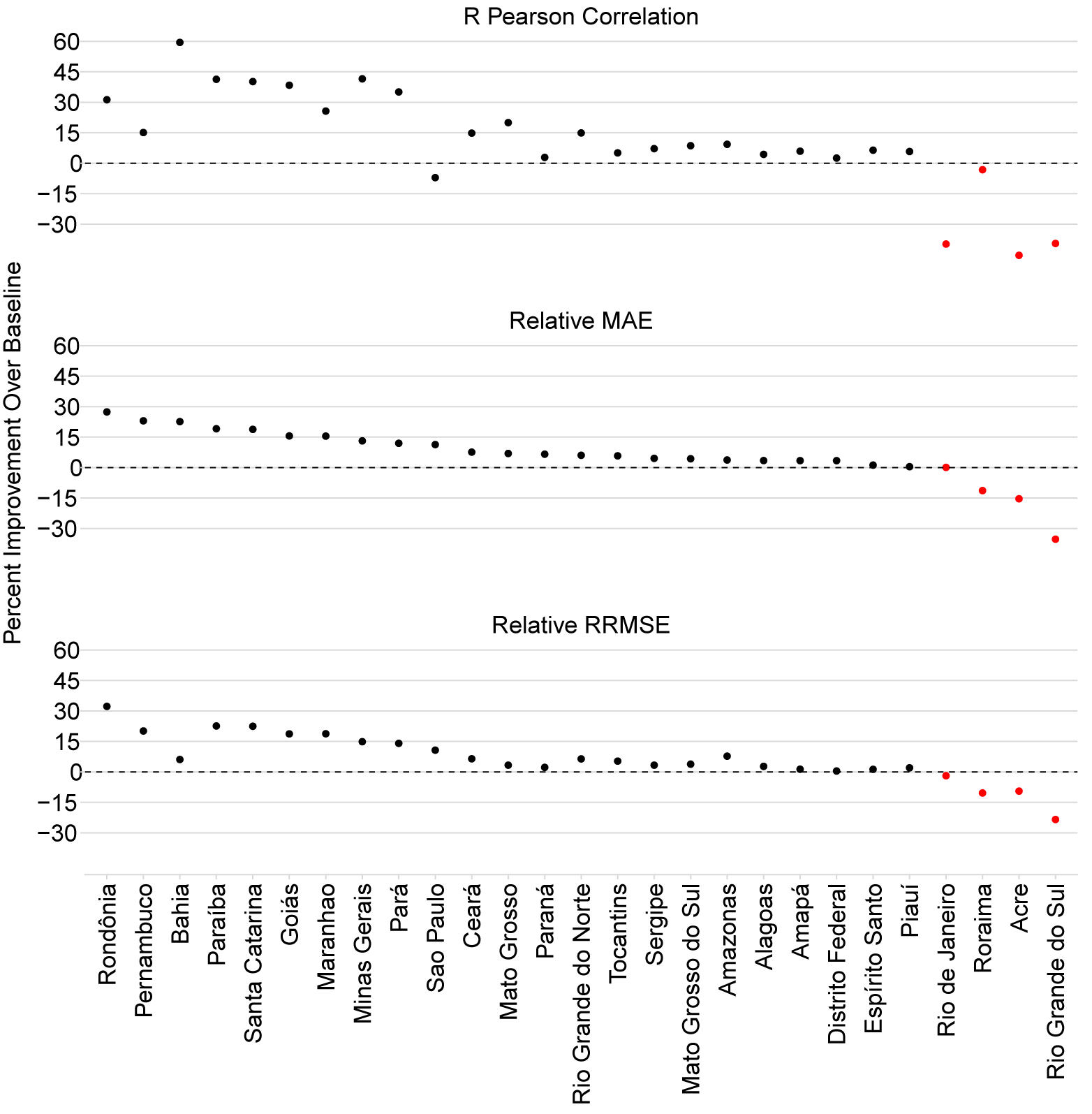
Relative improvement of nowcasting using exogenous data. The inclusion of exogenous data improved nowcasting over the baseline model (SARIMA or STL) in 22 or 23 states, depending on the error metric of interest. Red dots indicate states where the predictions were best on average using the STL model without exogenous data streams, so forcing the use of exogenous data made results worse.

**Figure 6.**
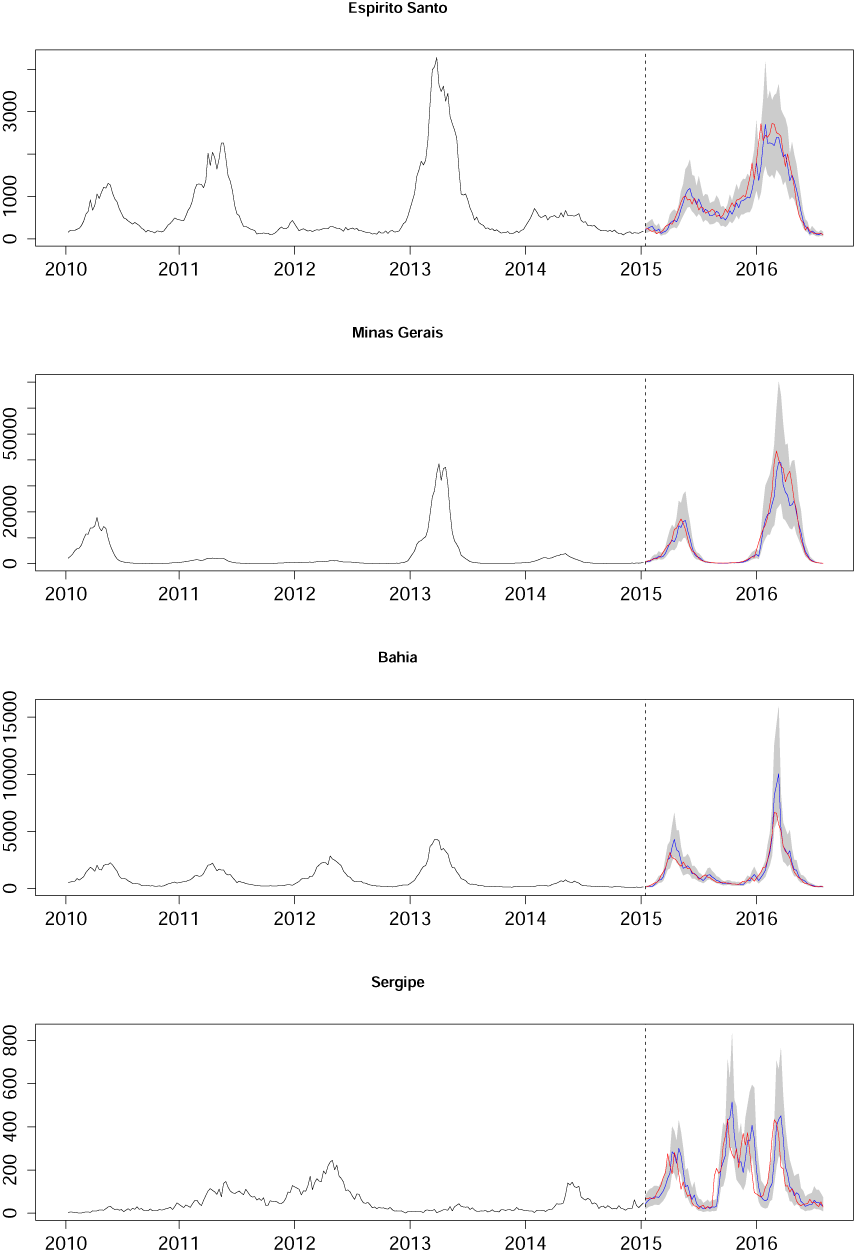
State-level observed vs. nowcasted dengue by highest ranked model. Time series of dengue are displayed for four Brazilian states. The vertical dashed line marks the boundary between training and testing weeks. The black time series corresponds to observed dengue in the training weeks (years 2010–2014). The blue line corresponds to predicted dengue; the red line corresponds to observed dengue within the testing weeks (years 2015–2016). The gray shadings are the associated 95% prediction intervals.

### Data sources, robustness of predictions, and patterns

As expected, data collection, data cleaning and data fusion (to align the time series) all proved challenging. We used standard kriging methods to fill in missing weather and satellite imagery data over the time series. The satellite imagery alone was over 30 terabytes; we derived indicators directly from the original satellite imagery rather than using “canned” products (which are typically used in the literature) so that we could improve resolution, include cloudy pixels as potential predictors, and have complete control over computation of indices, in turn allowing us to include some less common indices. There were also changes in municipality boundaries during the time period considered.

## Discussion

From this study, we first conclude that highly heterogeneous countries, such as Brazil, require data and analyses at localized geographical resolutions in order to correctly understand and capture the impact of complex interactions between environmental, population, and vector dynamics on dengue spread. That is, aggregating data at the national and even sometimes regional scale may provide misleading information about true dengue dynamics within an area. In fact, it is possible one could draw incorrect conclusions about risk and important predictors from aggregated data (e.g., Simpson’s paradox). Second, disparate data streams are necessary to model and nowcast dengue at various spatial scales and there is no single data stream that can be used for the entire country of Brazil. More data streams may also increase robustness of the model against loss of a single data stream in a “real-time” forecasting situation. Finally, our results show that using a variety of data streams (or a subset), along with several statistical prediction methods, can accurately predict dengue incidence in Brazil; this provides the foundation for a robust operational system for nowcasting and forecasting endemic dengue across broad geography at actionable scales.

In general, the scientific community has not taken full advantage of the big data revolution because of its limited ability to access and exploit real-time heterogeneous data streams in the context of computational epidemiology. We have developed a statistical data fusion approach that can systematically incorporate dynamic indicators to provide dengue risk and nowcast dengue using *volume* (i.e., around 30 terabytes of spatial and temporal data), *velocity* (i.e., weekly snapshots), *variety* (i.e., satellite, Internet, census, and climate), and *veracity* (i.e., curation of noisy and biased data).

As climate change increases the risk of mosquito-borne diseases like dengue across the globe, there is an urgent need for disease forecasting capabilities that can provide real-time decision support to reduce disease burden. Our results show that dengue dynamics can be accurately predicted with the exploitation of disparate, dynamic, and distributed data. However, there are several considerations in operationalizing this approach: 1) multiple data streams are needed to accurately nowcast dengue at multiple spatial resolutions especially for heterogeneous geographic environments, 2) multiple models are needed to obtain optimal results, and 3) the data collection and data fusion process must be streamlined (including accounting for sources that may go offline, change format, or miss values).

## Methods

We predict dengue at multiple temporal and spatial scales using different types of data sources: 1) long term (yearly) prediction of dengue risk maps using passive data (i.e., data that is not affected by dengue infections such as demographics data from the census, weather data, and satellite imagery) at the municipality, state, and country levels, 2) short term (weekly) dengue nowcasts (which is equivalent to two weeks ahead forecast assuming two weeks delay in the availability of case counts data) using both active (i.e., data sources that are affected by dengue infections such as Google Health Trends queries about dengue) and passive data (i.e., satellite imagery and weather) at the state level. Next, we briefly describe the data and methods used.

### Data

We collected five types of data for predicting dengue: dengue case counts, demographics, weather, satellite imagery, and Google Health Trends data. Features computed from all of these data sources are aligned spatially and temporally as follows. 1) The long-term (i.e., yearly) risk maps are predicted at the municipality level, which are then aggregated to predict dengue at the state and national level. Hence, for this model, all features are aligned at the municipality level and yearly scale. 2) With the short-term (i.e., weekly) nowcast, weekly dengue case counts are directly predicted at the state level and so for this model, features obtained from different data sources are aligned at the state level and weekly time scale.

### Dengue case counts

Data about dengue case counts were obtained from the Brazilian Ministry of Health from January 3, 2010 to July 17, 2017. These data are available for each of the 5564 municipalities in Brazil at the weekly time scale.

### Demographics

Demographics and socioeconomic data were obtained from the Brazil census at municipality and state levels for 2010. They include total 232 variables providing information about population statistics, education, income and poverty levels, and employment status.

### Weather

Daily temperature and humidity data were collected from weather stations in Brazil from April 1, 2009 to April 1, 2017 using the Global Surface Summary of the Day (GSOD) dataset from the National Oceanic and Atmospheric Administration (NOAA)^27^. There are 613 stations as compared to 5564 municipalities in Brazil, which means many of the municipalities are not directly covered by these weather stations. To address this, we used kriging methods to spatially interpolate temperature and humidity values.

### Satellite imagery

We used multispectral satellite images for each municipality in Brazil on a near-weekly basis from January 2010 to December 2016. These images came from four public-domain global coverage satellites (Landsat 5, Landsat 7, Landsat 8, and Sentinel-2), and were accessed using the Descartes Labs’ python-based platform^28^. Within each municipality, we computed the following indicators that provided information about factors related to dengue infections (e.g., water, vegetation): Normalized Difference Vegetation Index (NDVI), Green-based Normalized Difference Water Index (Green NDWI), Shortwave Infrared-based Normalized Difference Water Index (SWIR NDWI), Shortwave Infrared-based Normalized Burn Ratio (SWIR NBR), and the proportion of cloudy pixels.

For predicting risk maps, yearly weather and satellite-derived time series are obtained at the municipality level by computing a number of summary statistics on the features (minimum, maximum, mean, standard deviation, and range) by year, season (winter, spring, summer, and fall), and month (only for January and July, which are extreme in terms of dengue infections). This results in a total of 315 weather and 455 satellite features.

For predicting weekly dengue case counts, in addition to hourly temperature and humidity data, daily temperature range is also calculated. Then, both daily weather and satellite imagery data are converted to weekly features by computing summary statistics (minimum, maximum, and mean) for each municipality. Finally, these weekly municipality features are converted to state-level features by calculating another set of summary statistics (minimum, maximum, mean, and standard deviation) on them. This leads to a total of 40 weather feature time series and 52 satellite feature time series.

### Google Health Trends

Google is the most widely used search engine in Brazil with a market share of 97%^29^. It provides a normalized measure of region-specific search activities for the given terms through the Google Health Trends API^30^. We obtained GHT data at the country and state-level for Brazil from January 2011 to June 2017 on a weekly basis for the following dengue related terms (in both English and Portuguese): aedes, dengue, mosquito, mosquitoes, dengue virus, dengue fever, DHF, DENV, dengue hemorrhagic fever, aegypti, aedes aegypti, mosquito dengue, sintomas da dengue, aedes egípcio, egípcio, Vírus da dengue, novo vírus da dengue, Dengue é vírus, and dengue sintomas.

### Predicting Yearly Risk Maps

Our goal here is to predict yearly dengue risk maps using weather data, satellite imagery, and demographic data from the census. We used both normalized (using z score so that the mean is zero and the standard deviation is one) versions of raw values of features and log transformed values (computed as *log*(*x* + *x*_*min*_ + *σ*_*x*_) to deal with zero values and to account for differences in ranges of features). This leads to a large number of features (1000+) but some of them may be correlated, which may lead to unstable estimates of coefficients for regression analysis. To deal with multicollinearity, we first group together correlated features using agglomerative hierarchical clustering^31^ and then select representative features from each group.

In agglomerative hierarchical clustering, initially, each data point represents its own cluster and then two clusters with the smallest distance between them are merged together. This process is repeated until all data points belong to single cluster, and this process leads to a tree structure of clusters. As our goal is to group together correlated features, we used 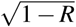 as a distance metric^32^, where *R* is a Pearson correlation between two clusters. We used single link criterion (i.e., distance between clusters *A* and *B* is defined as *min*{*d*(*a, b*)|*a* ∈ *A, b* ∈ *B*}) to merge two closest clusters as it guarantees that no two features with correlation above the threshold will be put in different clusters. Here, we represent each cluster by a set of representative variables. To select the optimal number of clusters, we perform multi-objective optimization which tries to minimize both the total number of representative variables selected for all of the clusters and the information lost due to keeping only these representative variables.

Suppose a cluster consists of *n* variables and we have selected *k* representative variables with indices *j*_1_, *j*_2_, …, *j*_*k*_ from it. Each of the remaining *n* − *k* variables can be eliminated and represented by one of the *k* representative variables. Information distance between a pair comprising eliminated variable *x*_*i*_ and representative variable *x* _*j*_ is defined as follows^33^:

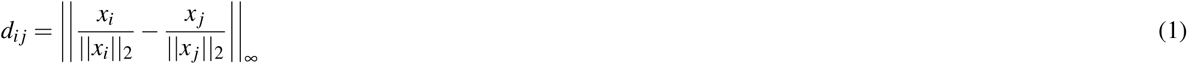

Each eliminated variable *x*_*i*_ can be represented by the representative variable *x* _*j*_ that minimizes the information distance between them as shown below:

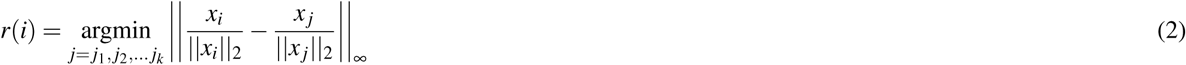

Here, *r*(*i*) is the representative variable for *x*_*i*_. The information loss for a cluster *c* consisting of *n* variables can be computed as

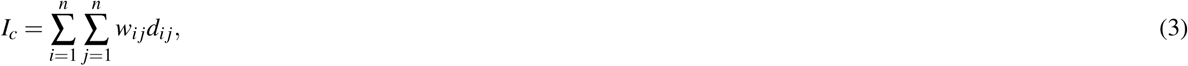

where *w*_*i j*_ = 1 if *r*(*i*) = *j* and 0 otherwise. Then the total information loss^33^ for a cluster assignment consisting of *C* clusters can be computed as

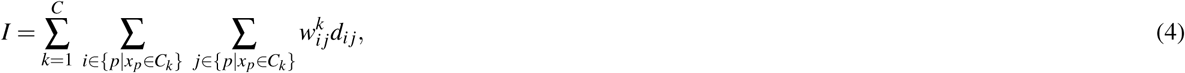

where *C*_*k*_ represents cluster *k*, and 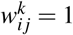 if variable *x*_*i*_ from cluster *k* is represented by variable *x* _*j*_ from the same cluster and 0 otherwise.

For multi-objective functions, there may be more than one non-dominated solution and the set of these non-dominated solutions is called the Pareto front. To obtain the Pareto front, we used NSGA-II^34^, one of the most widely used genetic algorithms with a simulated binary crossover operator^35^ and a polynomial mutation operator^36^ as recombination and mutation operators, respectively. The optimal solution (or point) in our case is one representative variable and zero information loss and so we chose the solution on the Pareto front that is at the minimum distance from this point.

Because the environmental variables (consisting of weather and satellite imagery variables) change over time but the census data (containing demographics) is static (as it is only available for 2010), clustering is performed separately for the environmental variables and demographics variables. The optimal clustering assignment selected 57 variables (with Pearson correlation threshold of 0.82) from the Census and 414 variables (with Pearson correlation threshold of 0.93) from weather and satellite imagery data sources. While this clustering approach selected a smaller set of relatively less correlated features, some of the selected features may not be very relevant in predicting dengue. Because of this, we used a generalized linear model (GLM)^37^ with forward and backward feature selection to predict the logarithm of yearly dengue case counts at the municipality level. Here, input to the GLM consisted of observations corresponding to municipality-year pairs. The predictions at the municipality level are aggregated to predict dengue case counts at the state and country levels. We tried both training one GLM model for the entire country and training separate GLM models for each of the macro regions, and region-specific models led to better performance.

### Predicting Weekly Dengue Case Counts

Our goal here is to nowcast dengue case counts (which is equivalent to two weeks ahead forecasts assuming two weeks delay in the availability of case counts data) at the state-level for Brazil. For this purpose, we explored a number of statistical models and heterogeneous data sources to identify the combination of statistical model and external data sources that lead to optimal out-of-sample predictions for each state. We explored the most common and well understood statistical time series models as they explicitly take into account temporal dependence and seasonal aspects of dengue infections and vary in inclusion of exogenous variables: Seasonal Trend decomposition using Loess (STL)^38^, Seasonal Autoregressive Integrated Moving Average (SARIMA) model^39^, SARIMAX with Principal Component Analysis (PCA), and SARIMAX with Partial Least Squares (PLS), and vector autoregression (VAR). The first two models (STL and SARIMA) only take into account past values of dengue case counts. On the other hand, the last models (SARIMAX with PCA, SARIMAX with PLS, and VAR with PCA) also consider exogenous variables from heterogeneous data sources (weather, satellite imagery, and Google Health Trends) and case counts from the neighboring states (to account for spatial dependency of dengue infections) as predictors. SARIMAX performed better than VAR across the board for exogenous variables, so we present SARIMAX results. To identify the set of heterogeneous data sources that are optimally predictive for each state, we tried each of these data sources individually, combinations of two, and all three of them together for each state for both SARIMAX with PCA and SARIMAX with PLS models. All SARIMAX with PCA and SARIMAX with PLS models include case counts from the neighboring states as predictors. Next, we briefly describe the statistical models used.

### Seasonal Trend decomposition using Loess (STL)

STL is a well known robust time series decomposition method that decomposes time series into seasonal, trend, and remainder components^38^. We tried both additive and multiplicative forms of STL method which decompose a time series as

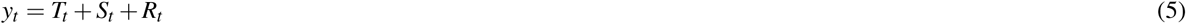

and

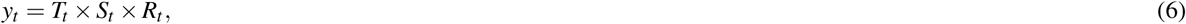

respectively. Here, *y*_*t*_ represents the response variable (i.e., case counts) at time *t* and *T*_*t*_, *S*_*t*_, and *R*_*t*_ represent trend, seasonal, and remainder components at time *t*. As the multiplicative form led to better predictions, we only present results from this model. Values of different parameters required to fit the STL model were chosen as follows. We assume periodicity of time series as 52 to reflect yearly dengue infection patterns, and the span of the loess window for seasonal and trend component extraction are set to 155 and 25, respectively, by comparing model fit within the training data set for a range of odd values (odd values are recommended by^38^). The seasonal component of the time series is estimated using loess smoothing and the de-seasonalized time series (trend and remainder components) are forecasted using the Autoregressive Integrated Moving Average (ARIMA) model (which is similar to the SARIMA model [described next] with no seasonal component).

### Seasonal Autoregressive Integrated Moving Average (SARIMA)

SARIMA is one of the most widely used time series forecasting models. It is denoted as *SARIMA*(*p, d, q*)(*P, D, Q*)_*m*_ where *p, d*, and *q* are autoregressive, difference, and moving average orders for the nonseasonal component, *P, D*, and *Q* are autoregressive, difference, and moving average orders for the seasonal component, and *m* is the number of observations per season (which is 52 weeks for our models)^39^. The SARIMA model is defined as

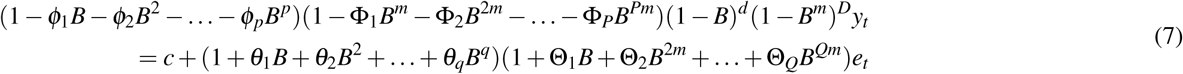

where *B* represents the backshift operator such that *By*_*t*_ = *y*_*t*−1_, *B*^2^*y*_*t*_ = *y*_*t*−2_, and so on; *ϕ* and Φ are coefficients for nonseasonal and seasonal autoregressive terms, respectively; and *θ* and Θ are coefficients for nonseasonal and seasonal moving average terms, respectively. Optimal values of *p, d, q, P, D*, and *Q* are chosen using the Akaike Information Criterion (AIC)^40^.

### Seasonal Autoregressive Integrated Moving Average with Exogenous Variables (SARIMAX)

SARIMAX model uses multiple linear regression based on exogenous variables to predict the response variable and then uses SARIMA model to predict the error from the multiple linear regression model. It is defined as follows:

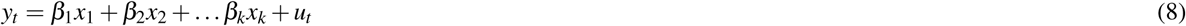

Here, *x*_1_, *x*_2_, … *x*_*k*_ are exogenous variables, *β* s are coefficients for these exogenous variables, and *u*_*t*_ is the error at time *t* which is modeled as

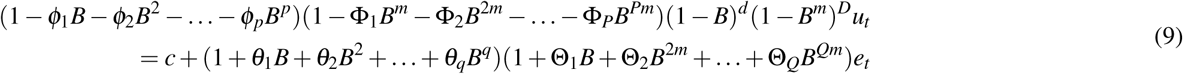

where all notation is similar to Equation 7.

Including information from weather, satellite imagery, and Google Health Trends data as well as case counts from the neighboring states leads to a large number of exogenous variables (100+). However, some of these variables may be correlated. Also, including a large number of predictors may lead to overfitting (i.e., reduced performance on new observations). To deal with these issues, we performed dimension reduction using two techniques: principal component analysis (PCA) and partial least squares (PLS). We also omitted any variables with zero variance. PCA is an unsupervised learning method that linearly transforms the data (exogenous variables in our study) into a new coordinate systems such that it maximizes the covariance of the data in lower dimension^41^. The direction of the largest variance is called the first principal component, the direction orthogonal to it and capturing the next largest variance is called the second principal component, and so on. As our goal is to predict dengue case counts, we tried all combinations of five PCA components (2^5^ = 32) that are most strongly correlated with dengue case counts as exogenous variables in SARIMAX models, and selected the one that led to optimal fit according to the AIC criterion. PLS can be viewed as a supervised version of PCA that also transforms the data such that it maximizes covariance of the predictors and response variables^42^. Similar to PCA, PLS components are also orthogonal but they are also highly correlated with the response variable. We tried all combinations of the top five PLS components (2^5^ = 32) as exogenous variables in the SARIMAX models and selected the one that led to optimal fit according to the AIC criterion.

For all of the models, we calculated point estimates and 95% prediction interval based on asymptotic normality of errors. In some rare cases, our models forecasted dengue case counts slightly below zero, which are converted to zero. We used R packages *pls*^43^ and *forecast*^44^ for PLS and training time series models, respectively.

### Model Evaluation

Statistical and machine learning models are prone to overfitting. Overfitting occurs when models are fitted to a particular training data set rather than to the underlying data generating phenomenon and hence they perform well on the training data set but fail to do well on new observations. To overcome the problem of overfitting, we used data from Jan 2010 to Dec 2014 as the training data and used data from Jan 2015 to July 2017 as test data for both yearly and weekly dengue predictions.

For predicting weekly dengue case counts, we assume two weeks of lag in availability of dengue case counts data and and produce two weeks ahead forecasts (e.g., data for weeks 1-261 is used to predict dengue case counts for week 263). We simulated a real world situation by adding one week of data to the training data set as it becomes available (i.e., data for weeks 1-261 is used to forecast week 263, data for weeks 1-262 is used to forecast week 264, and so on). When case counts from the neighboring states are included as predictors (for SARIMAX with PCA and SARIMA with PLS models), the two-week lag is addressed by forecasting case counts for neighboring states using SARIMA model. We assume that weather, satellite imagery, and Google Health Trends data is available in real time.

We used five metrics to measure the performance of the models: RMSE, RRMSE, Pearson correlation (R), MAE, and RMAE. These metrics are defined as follows:

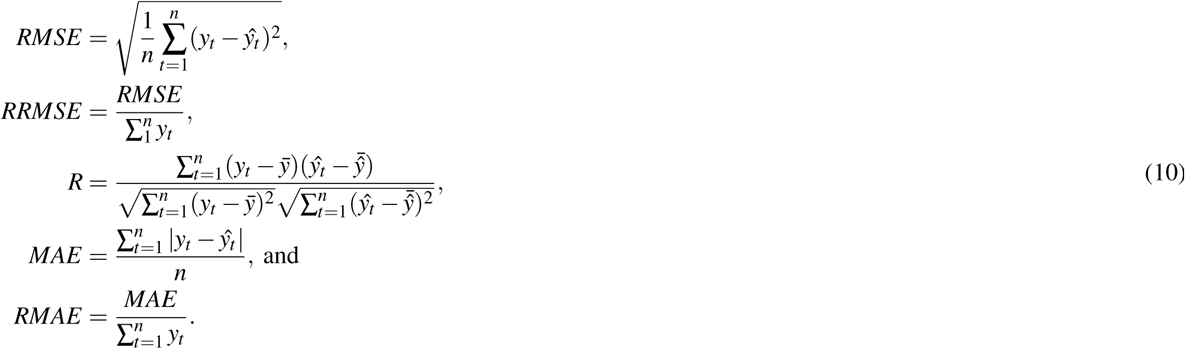

Here, *y*_*t*_ and *ŷ*_*t*_ are observed dengue case counts and predicted dengue case counts at time *t*, respectively and 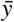 and 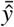 are mean observed case counts and mean predicted case counts, respectively. RMSE and MAE are used commonly to measure performance for numeric outcomes (e.g., output of regression analysis) but they are affected by total number of case counts and hence not comparable across different states. RRMSE and RMAE are their normalized versions that are comparable across different states, along with R. Importantly, these are also comparable across future studies with new or competing models.

## Acknowledgements

We acknowledge Descartes Labs for streamlined access to the satellite imagery and acknowledge the Ministry of Health of Brazil for reported case counts data. Research presented in this article was supported by the Laboratory Directed Research and Development program of Los Alamos National Laboratory, projects 20190581ECR and 20180740ER. The content is solely the responsibility of the authors and does not necessarily represent the official views of the sponsors. LC and SDV were supported in part by NIH/NIGMS grant R01-GM130668-02. The funders had no role in study design, data collection and analysis, decision to publish, or preparation of the manuscript. Los Alamos National Laboratory is operated by Triad National Security, LLC, for the National Nuclear Security Administration of U.S. Department of Energy (Contract No. 89233218CNA000001).

## Author contributions statement

CM co-conceived and coordinated the study, team, and methods, and helped write the manuscript; GF and AZ co-conceived the study, acquired, processed, and analyzed the satellite imagery and weather data, and helped write the manuscript; NP acquired GHT data and assisted with computational methods and data fusion; KK implemented the statistical methodology, assisted with figures, and helped write the manuscript; KM cleaned and fused data and assisted with figures; LC performed statistical analysis on spatial spread timing across states and assisted with figures; DO helped develop the statistical methodology and helped write the manuscript; AS performed kriging on the data; JC helped to plan the manuscript, fuse data, and analyze results; NG co-conceived the study and acquired case surveillance data; and SDV co-conceived the project, acquired and translated the census data, helped write the manuscript, and led the team. All authors reviewed the final manuscript.

## Additional information

The authors have no known competing interests.

## Data Availability Statement

The case counts data is available from the Brazilian Ministry of Health upon request; demographics data was from the publicly available Brazil 2010 census; weather data was obtained from Global Surface Summary of the Day (GSOD) dataset from the National Oceanic and Atmospheric Administration; satellite data was obtained from Descartes Labs using Landsat 5, Landsat 7, Landsat 8, and Sentinel-2 satellites; Google Health Trends data was obtained from through the Google Health Trends API which is publicly available. Data is available upon request from the corresponding author.

## Code Availability Statement

The code was developed in R with publicly available R packages.

## Notes

https://arxiv.org/abs/2006.02483

### Competing Interest Statement

The authors have declared no competing interest.

