## Supplemental Tables for "Unlocking the Predictive Power of Heterogeneous Data to Build an Operational Dengue Forecasting System"

| Yearly Risk Map Regional Models Error | | | | | | | |
|---|---|---|---|---|---|---|---|
| | | Municipality Level Error | | | State Level Error | | |
| | Year(s) | RRMSE | RMAE | R | RRMSE | RMAE | R |
| **Training** | **2010-2014** | **0.004646973** | **0.0001674771** | **0.14272417** | **0.10552409** | **0.02885302** | **0.7351028** |
| | 2010 | 0.012224943 | 0.0005552684 | 0.16009630 | 0.20008380 | 0.08272134 | 0.6947336 |
| | 2011 | 0.042747863 | 0.0012426705 | 0.07697522 | 0.75250462 | 0.22430254 | 0.7605392 |
| | 2012 | 0.009767673 | 0.0005710574 | 0.07880063 | 0.16620997 | 0.08714703 | 0.6993671 |
| | 2013 | 0.020112862 | 0.0011026660 | 0.21488345 | 0.58404851 | 0.20343547 | 0.8465791 |
| | 2014 | 0.008120369 | 0.0004353124 | 0.64814653 | 0.14594957 | 0.06319228 | 0.8883552 |
| **Testing** | **2015-2016** | **0.001152296** | **0.0001135361** | **0.71783585** | **0.02698789** | **0.01166679** | **0.9658939** |
| | 2015 | 0.002401753 | 0.0002352900 | 0.63897159 | 0.05984793 | 0.02514067 | 0.9672807 |
| | 2016 | 0.002133578 | 0.0002172325 | 0.80292967 | 0.04357084 | 0.02116986 | 0.9658786 |

| Yearly Risk Map State Models Error | | | | | | |
|---|---|---|---|---|---|---|
| | TrainTest | State_Name | UF | RRMSE_Error | RMAE_Error | R_Error |
| 1 | Test | Rondonia | 11 | 0.004669 | 0.002432 | 0.969833 |
| 2 | Train | Rondonia | 11 | 0.015705 | 0.005297 | 0.824937 |
| 3 | Test | Acre | 12 | 0.091738 | 0.017272 | 0.986758 |
| 4 | Train | Acre | 12 | 0.047116 | 0.010378 | 0.761996 |
| 5 | Test | Amazonas | 13 | 0.008037 | 0.001828 | 0.989536 |
| 6 | Train | Amazonas | 13 | 0.028402 | 0.002808 | 0.805497 |
| 7 | Test | Roraima | 14 | 0.013664 | 0.007472 | 0.981675 |
| 8 | Train | Roraima | 14 | 0.044211 | 0.016273 | 0.419314 |
| 9 | Test | Para | 15 | 0.003216 | 0.001148 | 0.959169 |
| 10 | Train | Para | 15 | 0.131669 | 0.009381 | 0.289206 |
| 11 | Test | Amapa | 16 | 0.015294 | 0.007426 | 0.966691 |
| 12 | Train | Amapa | 16 | 0.023667 | 0.010059 | 0.489205 |
| 13 | Test | Tocantins | 17 | 0.022537 | 0.002414 | 0.975098 |
| 14 | Train | Tocantins | 17 | 0.006168 | 0.001112 | 0.878823 |
| 15 | Test | Maranhao | 21 | 0.128941 | 0.011142 | 0.291558 |
| 16 | Train | Maranhao | 21 | 0.197339 | 0.010365 | 0.185700 |
| 17 | Test | Piaui | 22 | 0.019593 | 0.002144 | 0.983278 |
| 18 | Train | Piaui | 22 | 0.048903 | 0.002632 | 0.896314 |
| 19 | Test | Ceara | 23 | 0.029204 | 0.004131 | 0.334373 |
| 20 | Train | Ceara | 23 | 0.006834 | 0.000987 | 0.768624 |
| 21 | Test | Rio_Grande_do_Norte | 24 | 0.014474 | 0.003434 | 0.520109 |
| 22 | Train | Rio_Grande_do_Norte | 24 | 0.033400 | 0.002343 | 0.628321 |
| 23 | Test | Paraiba | 25 | 0.028346 | 0.004142 | 0.266589 |
| 24 | Train | Paraiba | 25 | 0.007011 | 0.000922 | 0.583519 |
| 25 | Test | Pernambuco | 26 | 0.018177 | 0.003980 | 0.467253 |
| 26 | Train | Pernambuco | 26 | 0.050548 | 0.003535 | 0.185611 |
| 27 | Test | Alagoas | 27 | 0.073379 | 0.011636 | 0.702762 |
| 28 | Train | Alagoas | 27 | 0.008544 | 0.001483 | 0.672988 |
| 29 | Test | Sergipe | 28 | 0.037708 | 0.008838 | 0.825473 |
| 30 | Train | Sergipe | 28 | 0.010092 | 0.002657 | 0.805042 |
| 31 | Test | Bahia | 29 | 0.009982 | 0.001647 | 0.879302 |
| 32 | Train | Bahia | 29 | 0.012581 | 0.001108 | 0.302736 |
| 33 | Test | Minas_Gerais | 31 | 0.005443 | 0.000580 | 0.881332 |
| 34 | Train | Minas_Gerais | 31 | 0.035890 | 0.001789 | 0.200809 |
| 35 | Test | Espirito_Santo | 32 | 0.069385 | 0.013729 | 0.749694 |
| 36 | Train | Espirito_Santo | 32 | 0.158130 | 0.030800 | 0.620589 |
| 37 | Test | Rio_de_Janeiro | 33 | 0.022973 | 0.006607 | 0.160777 |
| 38 | Train | Rio_de_Janeiro | 33 | 0.203715 | 0.019872 | 0.048986 |
| 39 | Test | Sao_Paulo | 35 | 0.006873 | 0.000889 | 0.771001 |
| 40 | Train | Sao_Paulo | 35 | 0.007565 | 0.000514 | 0.712931 |
| 41 | Test | Parana | 41 | 0.057973 | 0.005857 | 0.405073 |
| 42 | Train | Parana | 41 | 0.145616 | 0.006568 | 0.266662 |
| 43 | Test | Santa_Catarina | 42 | 0.138128 | 0.009206 | 0.102220 |
| 44 | Train | Santa_Catarina | 42 | 0.067988 | 0.003875 | 0.457490 |
| 45 | Test | Rio_Grande_do_Sul | 43 | 0.014579 | 0.001614 | 0.548884 |
| 46 | Train | Rio_Grande_do_Sul | 43 | 0.012428 | 0.000393 | 0.059978 |
| 47 | Test | Mato_Grosso_do_Sul | 50 | 0.002352 | 0.000950 | 0.995774 |
| 48 | Train | Mato_Grosso_do_Sul | 50 | 0.023893 | 0.003984 | 0.833053 |
| 49 | Test | Mato_Grosso | 51 | 0.001781 | 0.000748 | 0.975180 |
| 50 | Train | Mato_Grosso | 51 | 0.033101 | 0.003691 | 0.732382 |
| 51 | Test | Goias | 52 | 0.005062 | 0.000731 | 0.938787 |
| 52 | Train | Goias | 52 | 0.009818 | 0.001085 | 0.367924 |
| 53 | Test | Distrito_Federal | 53 | 0.174692 | 0.132744 | 1.000000 |
| 54 | Train | Distrito_Federal | 53 | 1.622634 | 1.038559 | 0.448355 |

| Model | | MAE | RMAE | R | RMSE | RRMSE |
|---|---|---|---|---|---|---|
| **SARIMA** | (Q0, Q2, Q4) | (6.20, 114.82, 2691.06) | (.002002, .00304, .008005) | (.678, .910, .956) | (8.70, 193.78, 5313.70) | (.00310, .00493, .0133) |
| | (Mean, SD) | (328.45, 601.70) | (.00348, .00127) | (.889, .0721) | (592.37, 1184.52) | (.00569, .00211) |
| **SARIMAX - PCA** | (Q0, Q2, Q4) | (6.23, 105.47, 2784.62) | (.00198, .00294, .00714) | (.625, .920, .975) | (8.75, **177.79**, 6225.67) | (.00311, .00471, .0146) |
| | (Mean, SD) | (316.54, 591.57) | (.00334, .00116) | (.891, .0795) | (618.68, 1288.59) | (.00576, .00256) |
| **SARIMAX - PLS** | (Q0, Q2, Q4) | (6.19, **104.16**, 2245.77) | (.00197, .00308, .00726) | (.618, **.923**, .970) | (8.69, 177.95, 4662.04) | (.000302, .00497, .0151) |
| | (Mean, SD) | (306.44, 525.27) | (.00336, .00116) | (.894, .0807) | (570.64, 1069.39) | (.00571, .00250) |
| **STL - add.** | (Q0, Q2, Q4) | (6.65, 121.11, 3164.44) | (.00287, .00409, .00807) | (.563, .871, .936) | (9.61, 201.68, 6197.06) | (.00390, .00667, .0133) |
| | (Mean, SD) | (421.28, 787.53) | (.00419, .00134) | (.842, .0920) | (728.32, 1441.13) | (.00683, .00258) |
| **STL - mult.** | (Q0, Q2, Q4) | (5.76, 126.07, 2532.55) | (.00241, .00370, .00717) | (.609, .891, .972) | (8.16, 205.97, 5221.25) | (.00369, .00616, .0108) |
| | (Mean, SD) | (363.97, 629.75) | (.00379, .00111) | (.876, .0807) | (683.30, 1302.39) | (.00647, .00205) |
| **VAR - PCA** | (Q0, Q2, Q4) | (7.65, 121.61, 3429.73) | (.00223, .00362, .00631) | (.716, .881, .944) | (10.35, 196.05, 5906.13) | (.00334, .0553, .00954) |
| | (Mean, SD) | (394.66, 769.71) | (.00387, .00109) | (.878, .0598) | (640.03, 1304.89) | (.00602, .00180) |
| **Ensemble - mean** | (Q0, Q2, Q4) | (5.94, 110.37, 2161.64) | (.00196, **.00275**, .00528) | (.733, .918, .964) | (8.36, 186.69, 4351.41) | (.00296, **.00459**, .00953) |
| | (Mean, SD) | (**291.03**, 510.72) | (**.00314**, .000901) | (**.906**, .0576) | (518.89, 995.13) | (**.00515**, .00159) |
| **Ensemble - w. mean** | (Q0, Q2, Q4) | (5.94, 111.95, 2114.24) | (.00198, **.00275**, .00543) | (.697, .918, .969) | (8.40, 185.30, 4114.09) | (.00290, .00460, .00876) |
| | (Mean, SD) | (291.63, 504.01) | (.00318, .000905) | (.904, .0637) | (**512.22**, 955.75) | (.00518, .00160) |

**Table 1. Summary of nowcast performance over Brazilian states.** For each model paired with performance metric, summary statistics over the 27 states are presented for the testing weeks (2015-16). Q0, Q2, and Q4 correspond to the minimum, median, and maximum, respectively.

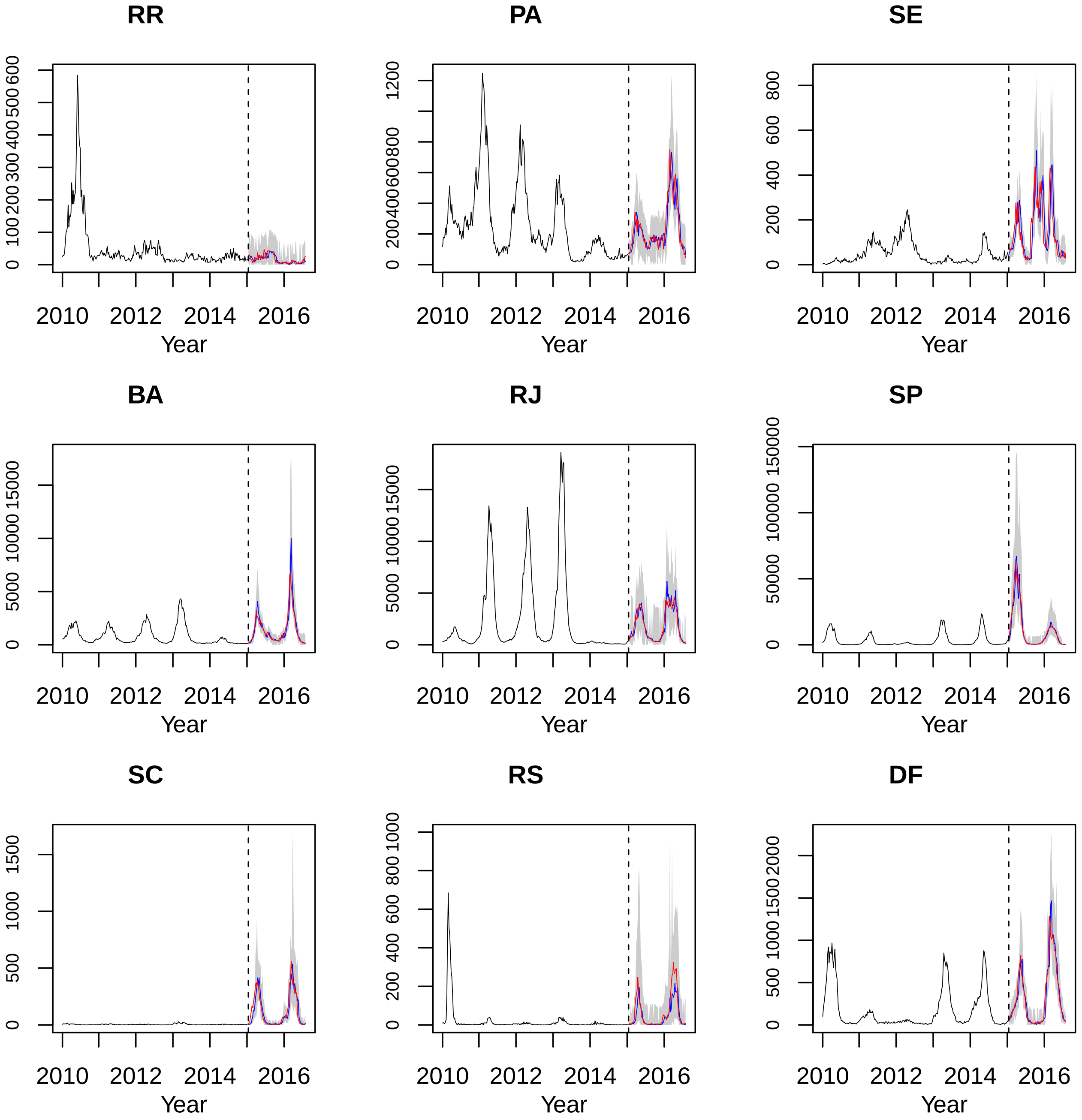


**Figure 1. Observed vs. nowcasted dengue by Brazilian state from trimmed mean ensemble approach.** Time series of dengue are displayed for a selection of Brazilian states. The vertical dashed line marks the boundary between training and testing weeks. The black time series corresponds to observed dengue in the training weeks (years 2010-14). The blue and red lines correspond to predicted and observed dengue, respectively, within the testing weeks (years 2015-16), while the gray shadings are the associated 95% prediction intervals. As displayed in Figure 1, the observed and nowcasted dengue counts for the testing weeks (2015-16) are very close for the selected states (results for all states are included as supplementary materials). In many cases, the observed dengue count is captured within the 95% prediction interval; empirical coverage for each state ranges from 81% to 100%, and the median and mean empirical coverage are approximately 96.3% and 95.3%, respectively. Our strong results show that dengue within a state can be nowcasted with high accuracy and confidence.

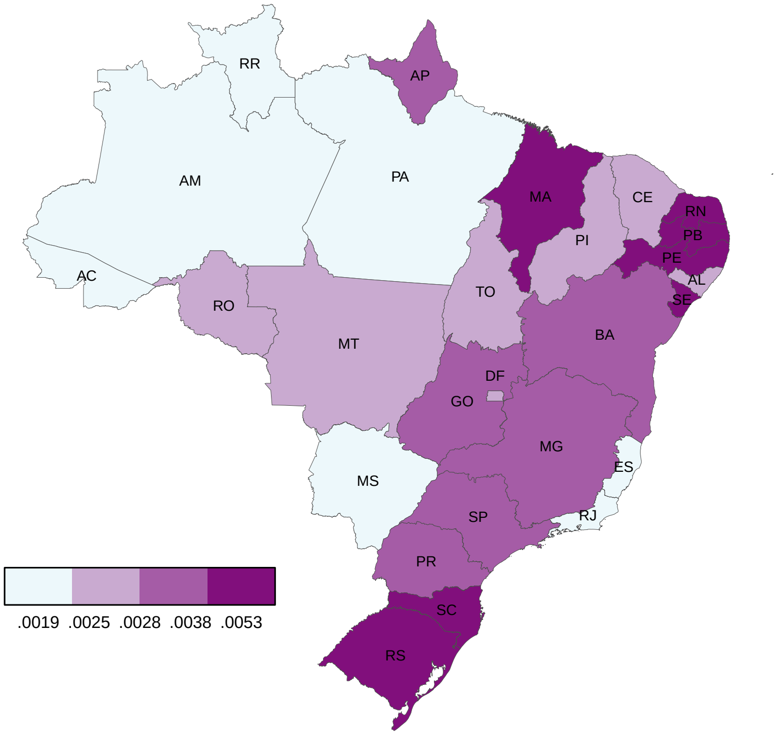


**Figure 2. Relative mean absolute error (RMAE) over the Brazilian states from the trimmed mean ensemble approach.** Among the 27 Brazilian states, we compare the testing RMAE, where map breaks are chosen as quartiles. There is a low but statistically significant spatial autocorrelation in the states' error.

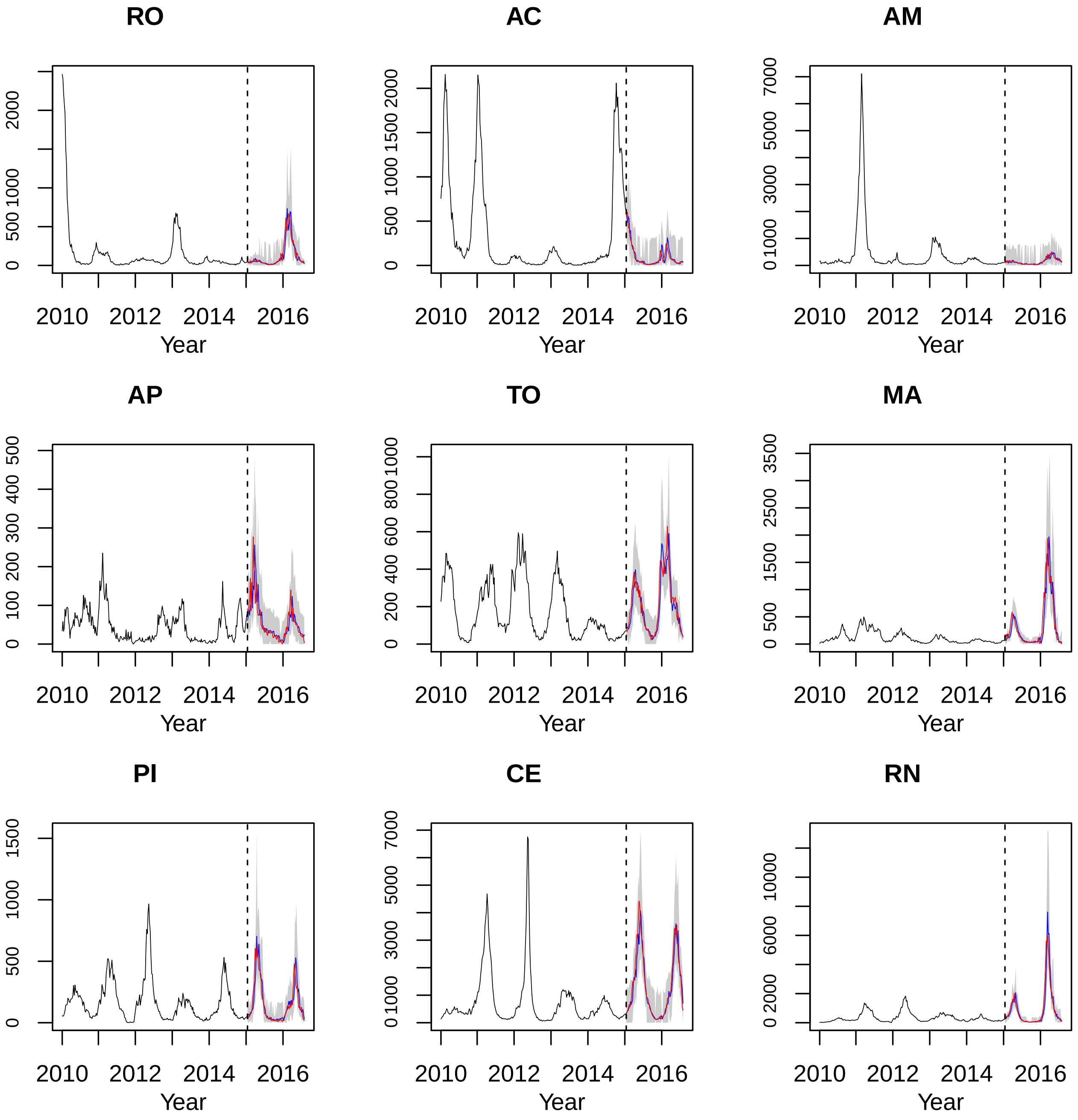


**Figure 3.** Observed vs. nowcasted dengue by Brazilian state from trimmed mean ensemble approach (2).

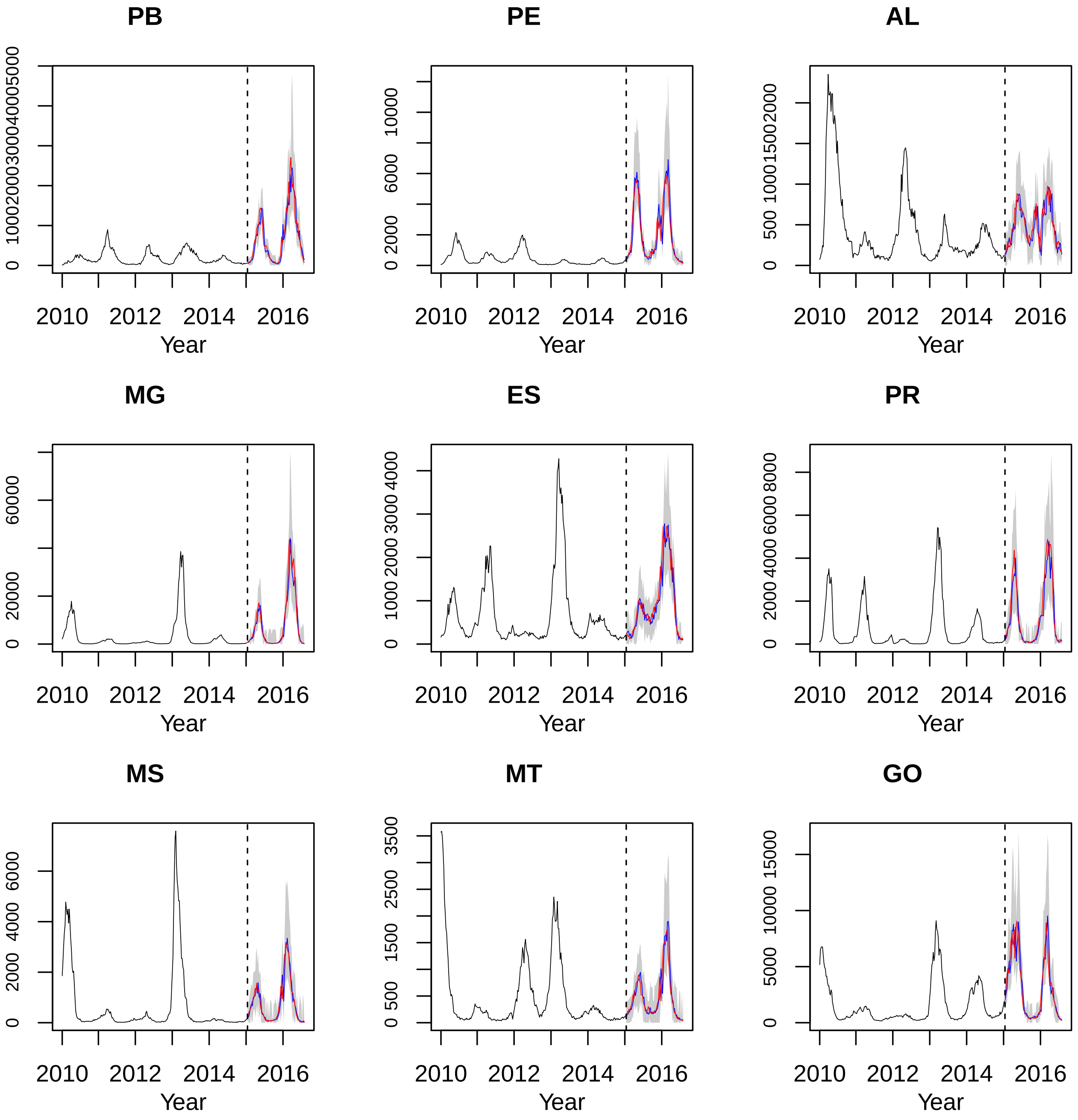


**Figure 4.** Observed vs. nowcasted dengue by Brazilian state from trimmed mean ensemble approach (3).
